# Age-related clonal haemopoiesis is associated with increased epigenetic age

**DOI:** 10.1101/600759

**Authors:** Neil A. Robertson, Ian J. Deary, Kristina Kirschner, Riccardo E. Marioni, Tamir Chandra

## Abstract

Age-related clonal haemopoiesis (ARCH) in healthy individuals was initially observed through an increased skewing in X chromosome inactivation. More recently, several groups reported that ARCH is driven by somatic mutations. The most prevalent ARCH mutations are in the DNMT3A and TET2 genes, previously described as drivers of myeloid malignancies. ARCH is associated with an increased risk for haematological cancers. ARCH also confers an increased risk for non-haematological diseases such as cardiovascular disease, atherosclerosis, and chronic ischemic heart failure, for which age is a main risk factor. Whether ARCH is linked to accelerated ageing has remained unexplored. The most accurate and commonly-used tools to measure age acceleration are epigenetic clocks. They are based on age-related methylation differences at specific CpG sites correlating chronological age accurately with epigenetic age. Deviations from chronological age towards an increased epigenetic age have been associated with increased risk of earlier mortality and age-related morbidities. Here we present evidence of accelerated epigenetic age in individuals with ARCH.

## Introduction

Age-related clonal haemopoiesis (ARCH) in healthy individuals was initially observed through an increased skewing in X chromosome inactivation ^1^. More recently, several groups reported that ARCH is driven by somatic mutations ^2,3^. The most prevalent ARCH mutations are in the DNMT3A and TET2 genes, previously described as drivers of myeloid malignancies. ARCH is associated with an increased risk for haematological cancers ^2,3^. ARCH also confers an increased risk for non-haematological diseases such as cardiovascular disease, atherosclerosis, and chronic ischemic heart failure, for which age is a main risk factor ^4,5^. Whether ARCH is linked to accelerated ageing has remained unexplored.

The most accurate and commonly-used tools to measure age acceleration are epigenetic clocks. They are based on age-related methylation differences at specific CpG sites ^6,7^, correlating chronological age accurately with epigenetic age. Deviations from chronological age towards an increased epigenetic age have been associated with increased risk of earlier mortality and age-related morbidities ^8,9^. Here we present evidence of accelerated epigenetic age in individuals with ARCH.

## Methods

### Study Cohorts

The Lothian Birth Cohorts (LBCs) of 1921 and 1936 are two longitudinal studies of ageing ^10^. Participants have been followed up every ~3 years, each for five waves, from the age of 70 (LBC1936) and 79 (LBC1921). Participants were community-dwelling, relatively healthy, and mostly lived in the City of Edinburgh or its surrounding area when recruited.

### Genome-wide methylation and sequencing

Whole blood DNA methylation levels were assessed using the Illumina HumanMethylation450 BeadChip. Quality control details have been reported previously ^11^. Genomic variants were determined in 1,148 LBC participants (n=873 and n=5 from waves 1 and 2 at mean ages 70 and 73 years, respectively in LBC1936; n=104 and n=166 at mean ages 79 and 87, respectively in LBC1921) with whole-genome sequencing (WGS) and methylation data. WGS data were aligned with Burrows-Wheeler Aligner and processed for non-unique or duplicate mapping reads with Picard Tools (genome coverage of 34.3 reads). Single-nucleotide variants and short indels were called with MuTect (v3.8) before annotation using the Ensembl Variant Effect Predictor alongside the Cosmic database of coding mutations (v86). ARCH variants were classified as per Jaiswal et al. ^2^.

### Epigenetic Age Acceleration

Epigenetic age acceleration was calculated online (https://dnamage.genetics.ucla.edu/home). We considered “BioAge4HAAdjAge’” and “AAHOAdjCellCounts’”, which are adaptations of the original Hannum and Horvath clocks that upweight for age-associated cell counts and control for white blood cell counts, respectively ^7^. Both epigenetic age estimates were regressed on chronological age to yield age acceleration residuals.

### Statistical Analysis

Linear regression adjusting for sex and methylation batch was used to determine the association between ARCH status (predictor) and Age Acceleration (response). All analyses were conducted in R v3.5.0.

## Results

Of the ten most prevalent ARCH mutations^2^, we had sufficient sample size and sequencing depth to annotate the top six in the LBC. We identified 71 participants (from 1,148) with ARCH (6.2%). The gene-specific prevalence ranged between 1-37 cases with ARCH-variant allele frequencies ranging from 0.034-0.677 (Fig.1A). Mutations in TET2 were exclusively frameshift and mutations detected in JAK2 (all V617F), SF3B1 and TP53 were exclusively missense.

**Figure 1.**
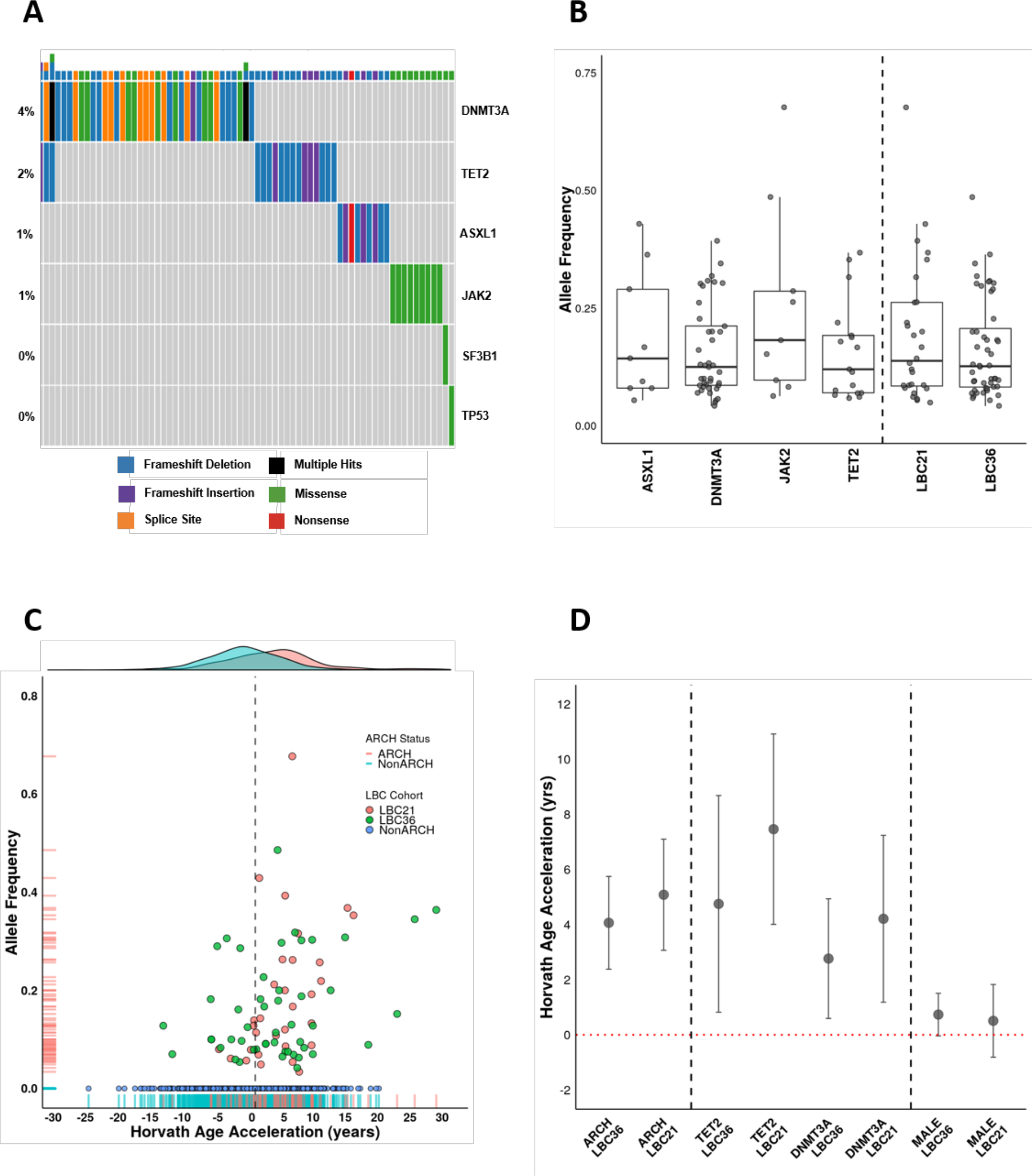
A) Oncoplot showing the type of variants within the ARCH positive subset of the Lothian Birth Cohort. This subset represents 71 participants (6.2% of 1,148 total) where one or more described somatic variants were detected in the six most prevalent ARCH-associated genes. B) Box and jitter plot describing the distribution of allele frequencies in all detected somatic ARCH variants. Genes with a single variant not shown are TP53 and SF3B1 (allele frequencies of 0.089 and 0.257, respectively). The overall distribution of allele frequencies by LBC cohort (LBC1921/LBC1936) is also displayed. C)Scatter plot showing the Horvath age acceleration (years) for individual LBC participants against the allele frequency of their somatic ARCH variant in both LBC1921 (orange dots, net 5.1 years; p=1.4×10^−6^) and LBC1936 (green dots, net 4.2 years; p=1.5×10^−6^) cohorts. Density plot highlighting the shift in distribution of Horvath age acceleration between ARCH positive (orange) and negative participant (turquoise) groups. D)Plot showing net age acceleration in ARCH (with 95% confidence intervals). The effect of sex (male versus female) on epigenetic ageing within the Lothian Birth Cohort is shown for comparison.

ARCH status was associated with a significant increase in Horvath’s epigenetic age acceleration: the increase was 4.2 (SE 0.9) years in LBC1936, and 5.1 (SE 1.0) years in LBC1921 (p=1.5×10^−6^ and 1.4×10^−6^, respectively) (Fig. 1C). Compared to non-ARCH carriers, those with TET2 mutations had a 4.8 (SE 2.0) year and 7.5 (SE 1.8) year increase in Horvath age acceleration in LBC1936 and LBC1921 (p=0.016 and p=3.2×10^−5^), respectively. Those with DNMT3A mutations had 2.9 (SE 1.1) years increase in LBC1936, and 4.2 (SE 1.6) years in LBC1921 (p=9.9×10^−3^ and p=6.9×10^−3^), respectively. (Fig.1D). These effect sizes are much larger than the sex differences in Horvath age acceleration, which were 0.75 (SE 0.4) years for men in LBC1936 (p=0.055), and 0.51 (SE 0.67) in LBC1921 (P=0.45) (Fig.1D). ARCH status was linked to increased Hannum age acceleration, albeit more modestly: 1.8 year (p=0.08) and 4.2 year (p=8.6×10^−3^) increase in LBC1936 and LBC1921, respectively (data not shown).

## Discussion

These results in the Lothian Birth Cohorts show a link between ARCH and age acceleration. This is an important conceptual advance leading to two, not necessarily mutually exclusive, testable scenarios: Firstly, ARCH could be an underlying cause for systemic ageing, explaining its link to various non-haematological, age-related diseases. Here, treating ARCH would have an effect on both, non-haematological and haematological disease. Secondly, an ageing microenvironment could enable clonal haemopoiesis, in which case non-haematological would be correlated, but not driven by ARCH. Here, treating ARCH would only affect haematological disease.

## Funding and Acknowledgements

The Lothian Birth Cohorts are funded by Age UK (Disconnected Mind grant). K.K. was supported by a grant from the NHSGG&C Research Endowment. T.C. and R.E.M. are supported by Chancellor’s Fellowships held at the University of Edinburgh. N.A.R. is supported by an MRC DTP studentship. R.E.M is supported by Alzheimer’s Research UK major project grant number ARUK-PG2017B-10. R.E.M. and I.J.D. are supported by The University of Edinburgh Centre for Cognitive Ageing and Cognitive Epidemiology (CCACE), part of the cross-council Lifelong Health and Wellbeing Initiative (MR/K026992/1); funding from the Biotechnology and Biological Sciences Research Council (BBSRC) and Medical Research Council (MRC) is gratefully acknowledged. The whole genome sequencing of the Lothian Birth Cohorts was funded through an award to the Roslin Institute from the Biotechnology and Biological Sciences Research Council (BBSRC).

